# Entropy Quantum Computing for Fixed-Backbone Protein Design

**DOI:** 10.64898/2026.02.20.706589

**Authors:** Babak Emami, Wesley Dyk, David Haycraft, Jenn Robinson, Lac Nguyen, Mohammad-Ali Miri, David J. Huggins

## Abstract

Computational protein design is a foundational challenge in biotechnology, advantageous for engineering novel enzymes and therapeutics, yet its combinatorial complexity remains a bottleneck for classical optimization. We formulate fixed–backbone computational protein design as a quadratic Hamiltonian over rotamer variables to naturally map onto a hybrid photonic entropy computing platform, Dirac-3. To assess solution quality and runtime performance, we benchmark the photonic solver against an exact classical cost function network (CFN) solver, which provides provably optimal baselines. For protein instances ranging from 493 to 943 variables, Dirac-3 attains solutions within 0.16–2.47 % of optimal energies. Empirical scaling analysis reveals a comparatively gentle effective runtime growth for the photonic solver over the measured regime, consistent with near-linear polynomial scaling, in contrast to the sharp super-polynomial growth observed for the classical baseline beyond approximately 1000 variables. These results suggest a near-term crossover regime in which hardware-aligned continuous-variable optimization may offer a practical promise for large computational protein design instances where exact classical methods become time-prohibitive.

## Introduction

Proteins are fundamental to biological systems, governing functions such as catalysis, cellular structure, and molecular recognition. Thus, the ability to design novel proteins with specified functions has enormous potential in drug discovery, synthetic biology, biomaterials, and enzyme engineering^1–5^. Fixed-backbone Computational Protein Design (CPD) is the task of designing amino acid sequences and their associated conformations that minimize an energy or free-energy model in the context of a fixed protein backbone, but unfortunately this task suffers from combinatorial explosion: for a protein with *N* positions and multiple rotamer choices per position, the number of possible sequence–conformation combinations grows exponentially. It can be shown that fixed-backbone CPD is an NP-hard problem^6^.

Classical methods have made progress in rendering CPD tractable for moderate size proteins. Traoré et al.^7^ reformulated fixed-backbone CPD as a Cost Function Network (CFN) and used the `toulbar2` solver to find the Global Minimum-Energy Conformation (GMEC) exactly for a range of mid-sized proteins. While extremely effective in terms of energy accuracy and runtime for many cases, these classical approaches can struggle as protein size, rotamer library size, or pairwise interaction complexity increase.

Concurrently, quantum and quantum-inspired optimization methods^8^ are being explored for molecular problems such as protein folding, docking, and molecular design^9^. Recent work includes QUBO formulations^10^ for peptide–protein docking^11^, variational quantum algorithms for coarse-grained protein folding^12^, and hybrid classical–quantum pipelines for drug discovery tasks^13^. These demonstrate promise but leave open questions about solution accuracy, scalability, and whether quantum devices can approach or outperform strong classical baselines in real CPD tasks.

In this work, we present a probabilistic formulation of fixed-backbone CPD mapped to Hamiltonians suitable for Quantum Computing Inc.’s entropy computing device, Dirac-3. We benchmark solution quality and runtime against the CFN baseline on a set of literature test-case proteins. Our results show that Dirac-3 can achieve energy values within 1-2 percent of the classical optimum in many cases. These findings demonstrate the viability of entropy-based optimization for high-dimensional sequence spaces and establish a scalable framework for integrating quantum-inspired hardware into practical protein engineering pipelines.

### Background and Related Work

The CPD problem has been studied for more than three decades, leading to the development of multiple algorithmic frameworks and software platforms^14–16^. These methods often focus on the fixed-backbone formulation, where the polypeptide backbone coordinates are held constant and the side-chain rotamer states are optimized to minimize an empirical force field. Despite its combinatorial complexity, fixed-backbone CPD has yielded biologically validated designs ranging from enzymes to binding proteins^17,18^.

Proteins are made from amino acids connected by peptide bonds and can be defined by their amino-acid sequence. The connected amino acids form a chain, termed the protein backbone, and different amino acids form branches with different protein sidechains. The backbone and sidechain of one amino acid is termed a residue (Figure 1a).

**Figure 1.**
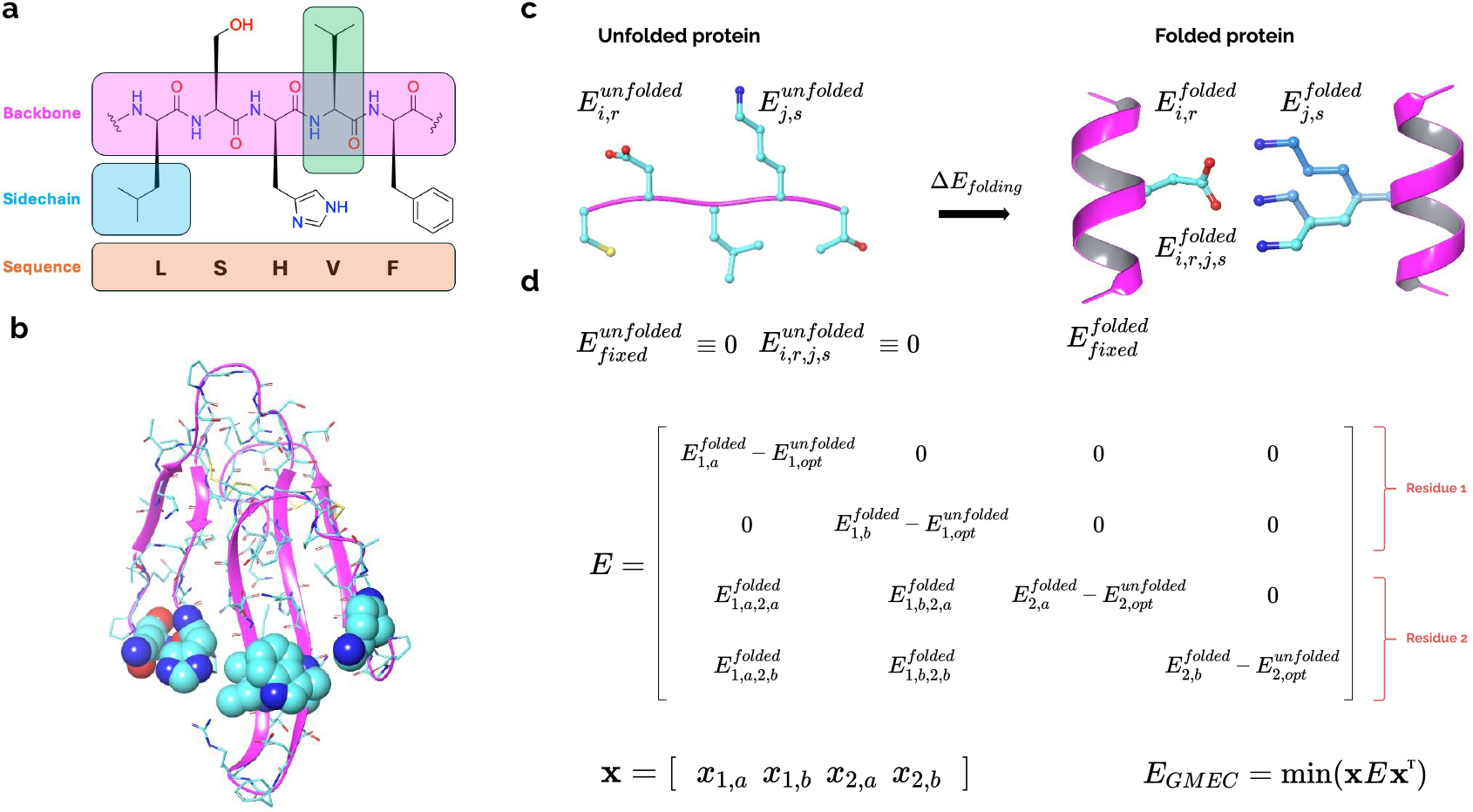
(a) Illustration of protein structure with backbone atoms boxed in magenta, a residue boxed in green, a sidechain boxed in cyan, and the sequence boxed in orange. (b) The folded structure of a protein, showing the backbone as a magenta ribbon and sidechains as cyan sticks. Five select sidechains are shown in cyan space filling to highlight the sidechain packing. (c) Illustration of the process of folding to yield the folding energy ΔE_folding_. The protein backbones are colored magenta and the sidechains are colored cyan. In the folded protein, three different rotamers of a residue are shown in shades of blue. The unfolded protein has an energy of zero for all fixed portions while the folded protein has a defined energy E_fixed_ for all fixed portions. The unfolded protein has energies E_i,r_ for each rotamer but the pairwise rotamer-rotamer interaction energies E_i,r,j,s_ are zero as they are far apart. The folded protein has energies E_i,r_ for each rotamer and E_i,r,j,s_ for each pair of rotamers. (d) An example of a simple energy matrix *E* for two residues, and a solution vector **x.** Solving to find the GMEC (and the optimal sequence in the case of CPD) is equivalent to minimizing **x**^*T*^ *E***x**.

For many proteins, the backbone folds into a distinct conformation such that the sidechains pack together to form intramolecular interactions (Figure 1b). Importantly, because proteins can be formed of hundreds or even thousands of residues, the 3D fold-space of a protein can be huge^19^. Determining the fold of a protein from its sequence is thus a very challenging problem^20^. Numerous approaches such as molecular dynamics have been applied to this problem, with success when applied to small proteins^21^. However, more recently, machine-learning approaches such as Alphafold^22^, Boltz^23^, and Chai^24^ have revolutionized the field of protein folding.

### The fixed-backbone approach using a rotamer library

In this work, we solve two problems directly related to protein folding: (1) identifying the global minimum energy conformation of a protein with a known fold and (2) identifying the amino acid sequence that minimizes the folding energy for a protein with a known fold. This approach has been studied many times previously^5^, typically using a pairwise additive forcefield^25^. For the purposes of solving this problem, the protein backbone is treated as being fixed and the sidechains for each residue can adopt a number of distinct conformations. These conformations are defined by a rotamer library and in this work we consider the widely used Dunbrack-Karplus rotamer library^26^. Each of the 20 natural amino acids is represented by a one-letter code or a three-letter code, and each can have multiple rotamers, which affect the pairwise interaction energies used to calculate the GMEC. These one- and three-letter codes and the rotamer numbers are included in the Supplementary Material (see Table S1). In fixed-backbone CPD, the protein backbone and any fixed sidechains have a defined energy (E_fixed_), each rotamer has a different interaction energy with the fixed protein backbone (E_i_), and each pair of rotamers has a different pairwise interaction energy (E_ij_).

The CPD problem is then suitable for optimization in this form, but as above it can lead to very large search spaces. As an example of the complexity: a five-residue protein with the sequence LSHVF from Figure 1a would have 6,561 possible combinations of rotamers using the Dunbrack-Karplus rotamer library. If one expands this to a CPD problem and allows each of the 20 natural amino acids at each position, there are now 27,675,731,591,168 possible combinations of rotamers.

We aim to calculate (i) the folding energy of a protein to its GMEC and (ii) for CPD the sequence yielding the most favorable folding energy. The folding energy is equal to the difference between the energy of the unfolded state and the folded state. For the unfolded state, the optimal (lowest energy) rotamer is selected at each position. For the non-fixed portions of the system, the energies for all rotamers at all positions can be formed into an energy matrix *E* for both the folded and unfolded states (Fig 1c). Any solution **x** can be represented as a binary column vector specifying which rotamer is selected for each residue (Figure 1c). The GMEC can be found by selecting **x** to minimize **x**^*T*^ *E***x** (Figure 1d) while respecting the constraint of selecting only one rotamer per residue. In this form, the problem is equivalent to the multiple-choice quadratic knapsack problem^27–29^ and has similarities to the Potts model as well as quadratic linearly constrained binary optimization (QLCBO). We aim to solve this problem using Dirac-3.

### Classical and quantum optimization approaches

Early CPD efforts relied on heuristic search strategies such as Monte Carlo sampling^30–32^ and dead-end elimination (DEE)^33–37^, which systematically prune rotamers incompatible with global optima^38,39^. These approaches, combined with stochastic minimization, powered widely adopted tools such as Rosetta^40^. More recently, exact optimization approaches have been developed^41^. Notably, Traoré et al.^7^ introduced a Cost Function Network (CFN) encoding of CPD and solved it exactly using the open-source solver `toulbar2`. Their results demonstrated that for mid-sized proteins, CFN can recover the GMEC efficiently, often within seconds^42^. Other classical frameworks include integer linear programming formulations^43–45^, branch-and-bound search, and graph-based decomposition techniques^46,47^. These methods achieve high accuracy but scale poorly with protein size and rotamer library expansion. Thus, while state-of-the-art classical solvers represent a rigorous baseline, the exponential scaling remains a fundamental limitation.

Along different lines, quantum and quantum-inspired methods have recently been proposed as alternatives for tackling CPD and related biomolecular problems. Perdomo-Ortiz et al.^48^ demonstrated that lattice-based protein folding models can be mapped to quadratic unconstrained binary optimization (QUBO) and solved using quantum annealers. Babbush et al.^49^ further developed adiabatic quantum simulations for protein folding, highlighting the potential of Hamiltonian-based encodings. More recently, variational quantum algorithms (VQAs) have been proposed for coarse-grained folding and docking tasks, exploiting hybrid classical–quantum pipelines to balance expressivity and near-term device constraints^12,13^. These studies suggest that quantum optimization devices may provide an advantage in navigating the high-dimensional energy landscapes of biomolecules, though large-scale demonstrations remain elusive.

Entropy computing was recently introduced as a quantum optimization paradigm that minimizes Hamiltonians within an open photonic quantum system that utilizes photon number state encoding and Fock basis readouts^50^. While such a paradigm can enable quantum-parallel computation^51^, its intermediate hybrid implementation through electronic measurement-and-feedback demonstrated efficient optimization with all-to-all graph connectivity and a high dynamic range^50,52^. This allows the native encoding of dense, multi-scale protein interaction graphs without the need for rescaling or complex embedding. The Quantum Computing Inc.’s entropy computing device, Dirac-3, is described in greater details in the Supplementary Material.

## Results

### Problem Formulation

Formulating the CPD problem for solution using Dirac-3 requires expressing the problem as a polynomial optimization objective, ideally up to quadratic terms. CPD aims to identify amino acid sequences and side-chain conformations that minimize a molecular energy function for a predefined backbone scaffold. This requires balancing the combinatorial search over residue identities and rotameric side-chain states with the evaluation of energetic contributions such as van der Waals forces, electrostatics, solvation, and hydrogen bonding^17,40,53^. Formally, the problem can be expressed as a discrete optimization task in which each residue position is associated with a catalogue of candidate rotamers, and the objective is to select exactly one rotamer per position so as to minimize the total energy. The highly-constrained nature of the CPD problem makes it a challenging optimization problem. Let *N* denote the number of residue positions in the fixed protein backbone, and let *R*_*i*_ represent the set of all feasible rotamers at position *i*. For each pair (*i, r*) we introduce a probability variable *x*_*i,r*_, where 0 ≤ *x*_*i,r*_ ≤ 1 for *i* ∈ *{*1,…, *N}* and *r* ∈ *R*_*i*_. Here *x*_*i,r*_ indicates the probability of selecting rotamer *r* at position *i*. Since exactly one rotamer must be chosen at each site, we impose the normalization constraint 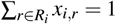 for all *i* = 1, …, *N*. Then, the energy of a candidate design is modeled as a sum of single-body and pairwise terms, where, single-body energies *E*(*i, r*) capture internal rotamer strain and side-chain–backbone interactions, while pairwise energies *E*(*i, r*; *j, s*) represent the physical interaction between rotamer *r* at position *i* and rotamer *s* at position *j*.

This classical decomposition is standard in CPD force fields and underpins leading design platforms such as Rosetta^17,40,54^ and OSPREY^45^. Accordingly, the optimization problem is cast as

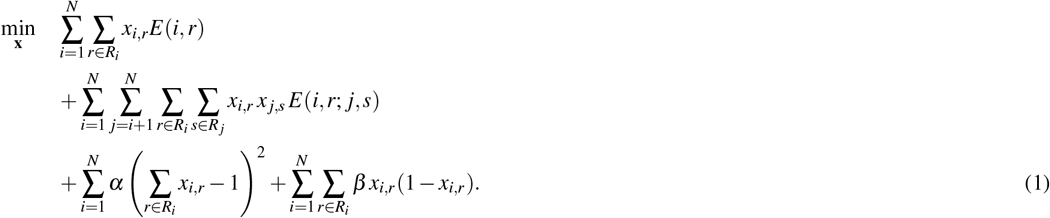

Here the quadratic penalty terms, controlled by hyper-parameters *α* and *β*, enforce discrete feasibility by discouraging violation of normalization and integrality. This relaxation allows us to map the CPD problem into a continuous optimization framework, while preserving exact recovery of discrete solutions for asymptotically large *α* and *β* . Finally, since *N* positions must each be assigned exactly one rotamer, we enforce the global constraint

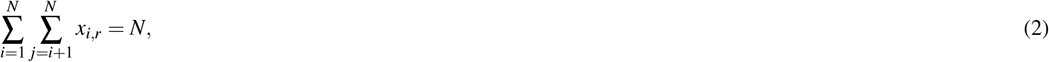

to ensure proper normalization across the entire problem.

This formulation yields a quadratic constrained optimization problem with both linear and quadratic terms, naturally expressible in Hamiltonian form for entropy-based optimization on QCi’s Dirac-3 device.

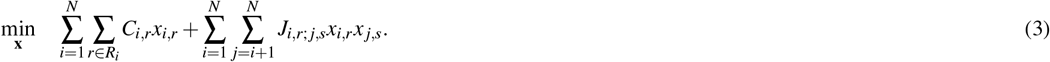

In this relation, the linear bias for rotamer (*i, r*) is given by *C*_*i,r*_ = *E*(*i, r*) − 2α + β where, *E*(*i, r*) is the single-body contribution, and *α, β* are the multipliers associated with normalization and integrality penalties, respectively. In addition, pairwise couplings are _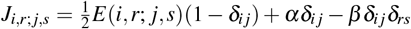_, embedding both rotamer interactions and feasibility constraints within the Hamiltonian.

Such a mapping parallel those used in QUBO formulations of CPD and related lattice protein folding problems^48, 49^. The advantage of this encoding is that it captures both classical CPD constraints and the structure needed for Hamiltonian-based solvers.

### Subproblem Formulation For Larger Test Cases

While mid-sized protein-design instances (up to 953 variables) can be mapped and solved directly on Dirac-3, larger systems entail thousands of rotamer states which makes obtaining a solution on Dirac-3 impractical. For two larger test cases, we therefore adopt a divide-and-conquer approach based on graph partitioning: we compress the full rotamer interaction into a position-level graph and split that graph into *k* balanced, weakly interacting blocks. Each block yields a sub-Hamiltonian that fits on Dirac-3; a solution is then constructed iteratively. This strategy preserves strong intra-block interactions while minimizing cross-block couplings, which in CPD often reflect weaker long-range effects. This decomposition approach is detailed in the Methods Section.

### Optimization Results

We evaluate the proposed Dirac-3 workflow on a canonical fixed-backbone CPD benchmark suite spanning small-sized proteins (1MJC, 1CSK, 1SHF, 1SHG, 1NXB, 1TEN, 1POH) and two larger instances (1RIS, 1GVP). We compare against the GMEC from the classical CFN baseline implemented in `toulbar2`, a strong reference for weighted-constraint encodings of CPD^7,55,56^. For the larger systems we apply the graph-partitioning framework described in Methods Section, solving sub-problems on Dirac-3.

Table S2 of the Supplementary Material reports problem sizes and coefficient dynamic ranges for all of the proteins studies in this paper. The dynamic range, defined as the ratio between the largest and smallest coefficients in the Hamiltonian (expressed in decibels), significantly influences numerical stability and optimization difficulty. Notably, Dirac-3 has a known performance limitation at 23 dB, making it particularly relevant to evaluate how it performs when handling Hamiltonians that exceed this range. The GMEC is known for each of these test cases, allowing effective benchmarking.

Figure 2 shows the solution energies for small protein problems, sorted by problem size, obtained by Dirac-3 and in comparison with the known optimal solutions from CFN^7^. For each problem, the results were collected from 100 runs. Dirac-3 finds solution energies within 0.16–2.47% of the GMEC, with a mean absolute gap of 1.21% and median 1.08%. The discrepancy between Dirac-3 and CFN energies can be better seen in Table 1.

**Table 1.** Energy and runtime comparison against the CFN baseline.

| Problem | Size | Avg. Sample Runtime (s) | CFN Runtime (s) | Best Dirac-3 Energy | CFN Energy |
| --- | --- | --- | --- | --- | --- |
| 1MJC | 493 | 4.23 | 0.08 | -70.45 | -70.71 |
| 1CSK | 616 | 3.54 | 0.15 | -55.41 | -55.50 |
| 1SHF | 638 | 4.38 | 0.17 | -60.07 | -61.60 |
| 1SHG | 737 | 9.38 | 0.25 | 65.13 | -65.56 |
| 1NXB | 800 | 3.86 | 0.24 | -37.35 | -38.27 |
| 1TEN | 808 | 6.11 | 0.33 | -91.45 | -92.45 |
| 1POH | 943 | 17.45 | 0.45 | -97.53 | -98.78 |

**Figure 2.**
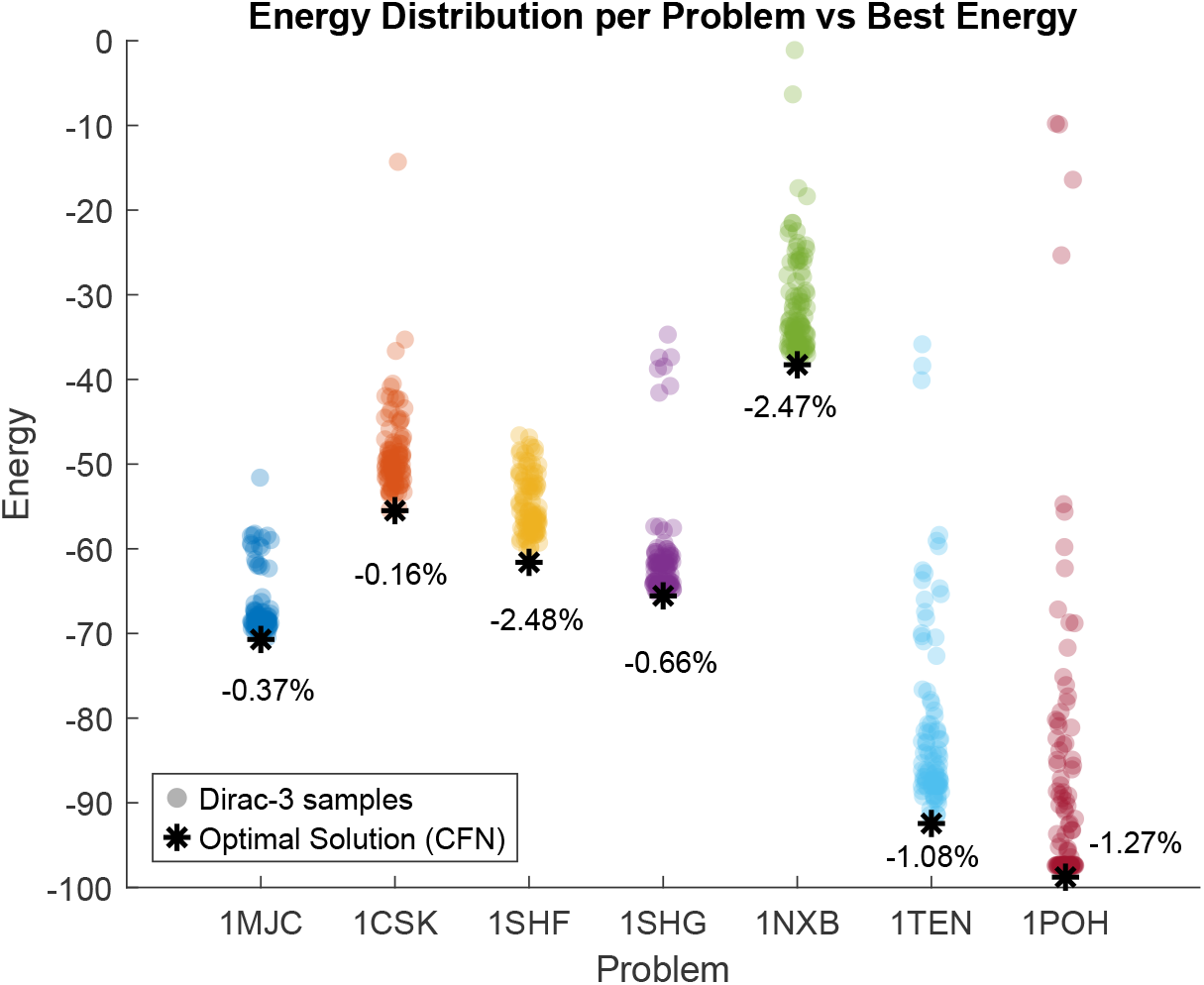
Energy distribution across 100 samples of Dirac-3 on each tested protein problems (circle), in comparison with the optimal energies from CFN^7^ (star marker). For each protein, Dirac-3 results correspond to 100 samples.

We next explore the effect of hardware parameters and mapping hyper-parameters. The goal is to identify robust operating regimes that retain energy quality while minimizing runtime. In what follows, the results of a thorough hyper-parameter study on the benchmark problem 1MJC is presented.

The *mean photon number* is an important hardware-level parameter on Dirac-3 that controls the optical intensity used to encode Hamiltonian coefficients within the photonic subsystem. Intuitively, it sets the average number of photons present in the system during computation, which in turn determines the signal-to-noise ratio and the precision with which energy differences can be resolved. In general, low mean photon numbers yield high-fidelity energy landscapes but longer runtimes due to weaker optical signals, whereas higher mean photon numbers accelerate convergence by strengthening signal amplitudes but risk saturating the device or introducing noise if pushed too far. Relaxation schedule is another important hardware parameter. In the context of QCi’s Dirac-3 device, *relaxation schedule* refers to the sequence of entropy-driven updates applied during the minimization process, effectively controlling how gradually the system relaxes from an initial high-entropy state to a low-entropy configuration that encodes a solution. In general, shallow schedules (few steps) yield faster convergence but risk trapping in local minima, while deeper schedules (more steps) provide improved energy accuracy at the cost of longer runtimes. The sample solution energies and their distributions at various values of relaxation schedule and mean photon number are shown in Figure 3(a) and (b), respectively. The corresponding runtimes and solution energies are shown in Tables S3 and S4 of the Supplementary Material. We observe that using a mean photon number of 0.003 seems to be optimal; using smaller mean photon numbers only increases runtime without any energy gain. We also observe that increasing schedule depth from 1 to 4 monotonically increases runtime without material energy gains beyond schedule 2.

**Figure 3.**
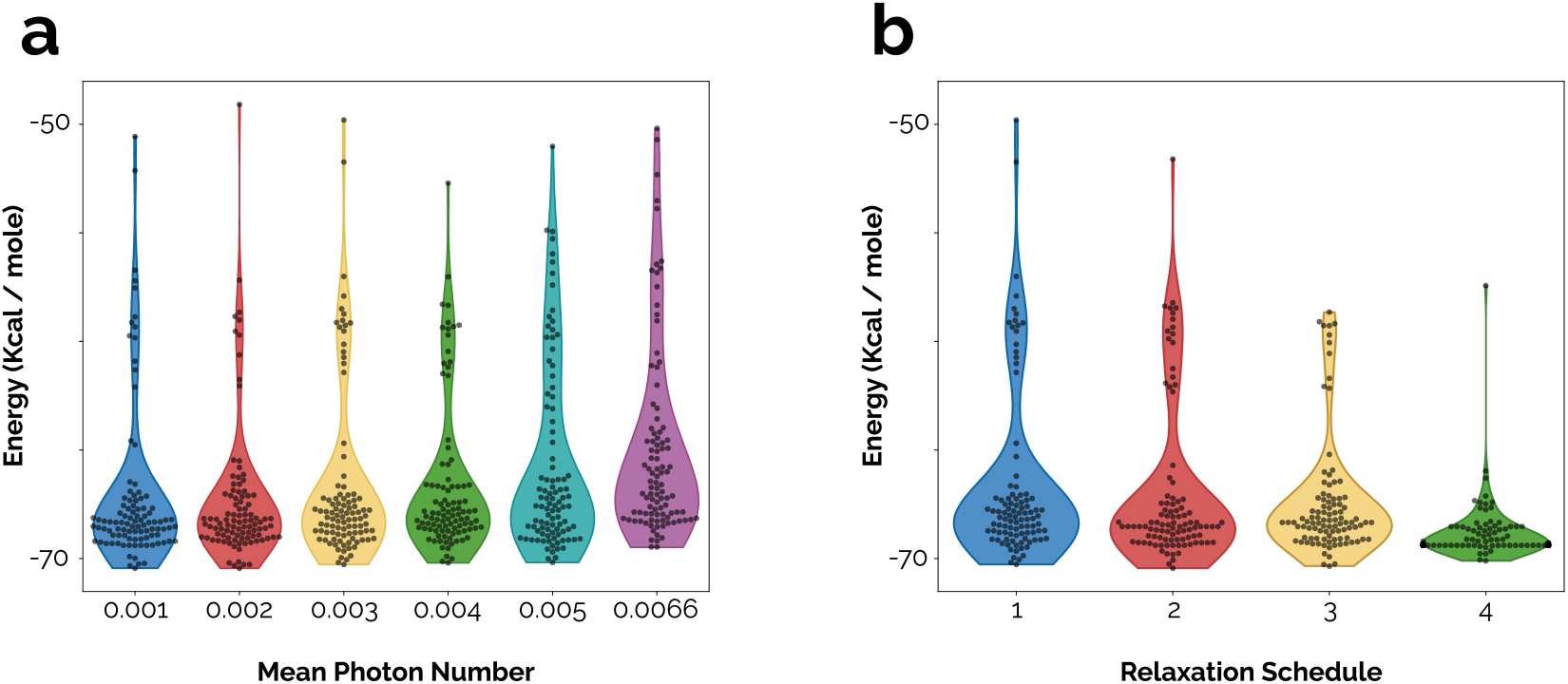
Impact of (a) mean photon and (b) relaxation schedule on solution energy for 1MJC protein.

Next, we investigated the impact of the dynamic range on the solution qualities. It is generally expected that pruning very large and very small coefficients helps reduce dynamic range, but aggressive pruning risks losing important interactions. The results are summarized in Tables S5 and S6 of the Supplementary Material. We observe that max-thresholds in the 5–10 range yield better solution energies by altering the Hamiltonian just enough to keep the dynamic range manageable; pushing to 100 degrades solution energy due to larger dynamic range values. In addition, min-thresholds around 10^−4^ to 10^−3^ are conservative and stable; while 10^−1^ happens to give a slightly lower energy here, that behavior is instance-specific and risks over-pruning. We have also studied the impact of penalty multipliers *α* and *β* (see equation 1) as shown in Tables S7 and S8 of the Supplementary Material.

Similar hyper-parameter study done on the other small-sized proteins studied here, namely 1CSK, 1SHF, 1SHG, 1NXB, 1TEN, and 1POH, yield similar results.

### Large Proteins

For the larger benchmarks in this study, namely 1RIS and 1GVP, the number of variables is much greater than the Dirac-3 limit of 953 variables. We used the approach discussed in Methods, by creating a position graph for each protein and partitioning it using the METIS algorithm. This partitions the full problem into sub-problems that can be solved on Dirac-3. Figure 4 visualizes the position graphs for the 1RIS and 1GVP proteins. It is worth noting that each node in these graphs represent a position on the protein backbone, however, the spatial position of graph nodes do not correspond to the actual spatial position in the molecular structure. Note also that the stronger interactions are shown by thicker graph edges. The partitions computed by METIS algorithm are color-coded in each graph. The partition counts are chosen such that the sub-problems are around 500 variables large. In both cases iterating twice over the sub-problems converged to steady solutions. Figure 4 also depicts the distributions of Dirac-3 sample energies obtained for 1RIS and 1GVP. The solution energies are compared to the corresponding CFN benchmarks in Table 2. For the two larger proteins (1RIS and 1GVP) solved via partitioning, the gap is around 7%. This is because the partitioned workflow is inherently heuristic: it optimizes a sequence of coupled subproblems and can therefore converge to a consistent low-energy configuration without guaranteeing recovery of the global minimum-energy conformation (GMEC).

**Table 2.** Energy and runtime comparison against the CFN baseline.

| Problem | Size | Avg. Sample Runtime (s) | CFN Runtime (s) | Best Dirac-3 Energy | CFN Energy |
| --- | --- | --- | --- | --- | --- |
| 1RIS | 3276 | 55.35 | 501 | -127.17 | -137.48 |
| 1GVP | 3826 | 65.42 | 594 | -114.48 | -123.56 |

**Figure 4.**
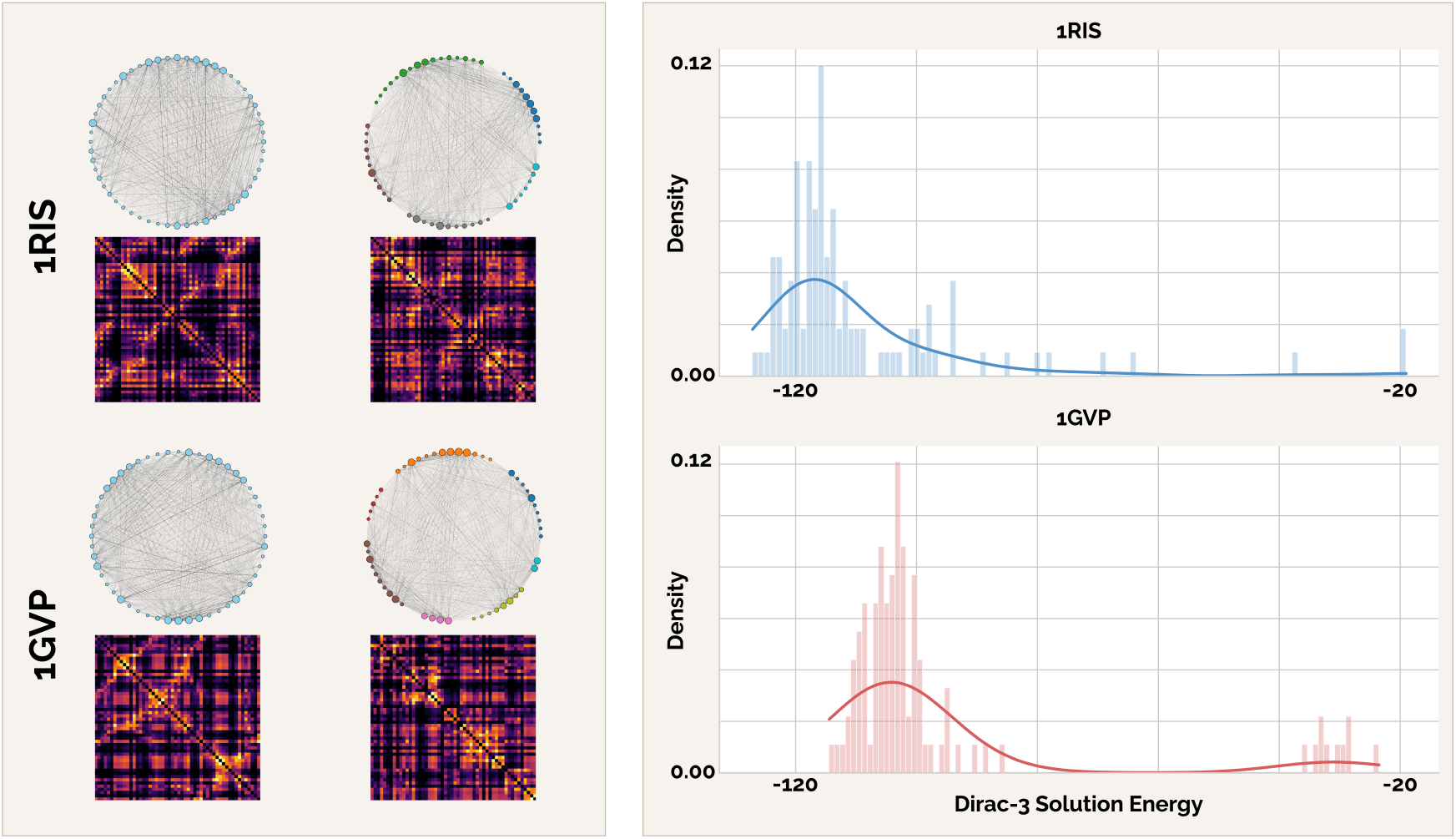
Solution of larger proteins 1RIS and 1GVP on Dirac-3 using the partitioning method; graph partitionings (left), and solution energy distributions (right).

### Runtime comparison and scaling behavior

Figure 5 compares wall-clock runtimes for the classical exact CFN solver and the Dirac-3 workflow as a function of problem size (number of rotamer variables). The plot includes observed Dirac-3 runtimes over the directly-solved regime (up to 943 variables). Over this range, Dirac-3 exhibits only mild growth in runtime with problem size, consistent with an effectively low-order scaling dominated by device-level overheads and fixed schedule costs rather than strongly size-dependent combinatorial search.

**Figure 5.**
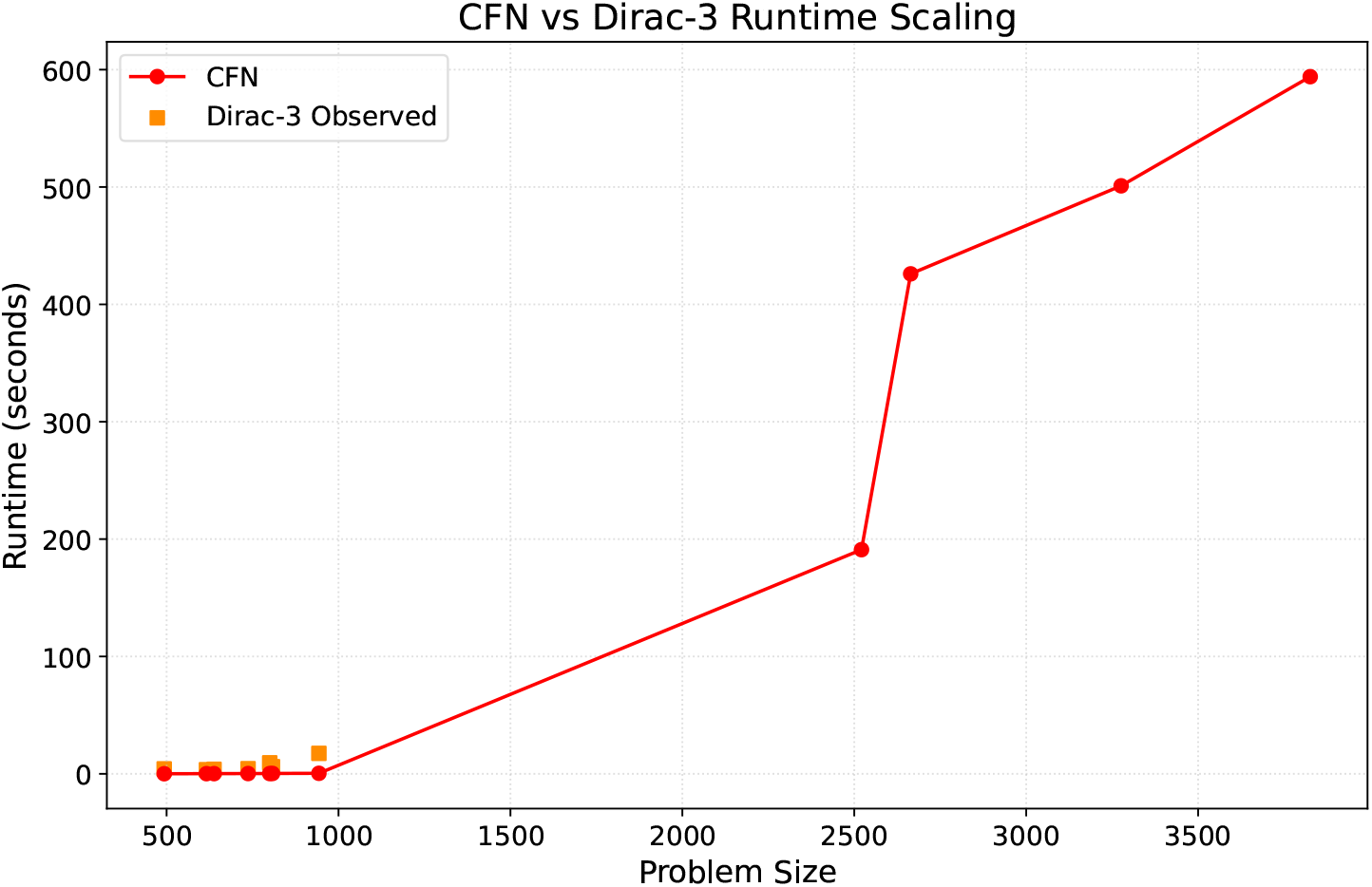
Comparison of runtime scaling between the classical exact CFN solver and the Dirac-3 on protein design problems.

Although CFN is extremely fast for the smallest instances, its runtime increases sharply with problem size, and the growth becomes pronounced beyond roughly the ∼ 1000-variable scale, consistent with the expected worst-case behavior of exact discrete optimization. The empirical separation between the curves suggests a practical regime in which entropy quantum optimization can potentially offer substantial wall-clock advantages once exact classical runtimes begin to escalate. The observed divergence between CFN and Dirac-3 trends is consistent with a near-term crossover region (roughly 1, 000–2, 000 variables) in which the runtime cost of exact classical optimization grows significantly while the Dirac-3 workflow remains operationally efficient. Further work will be required to test the scaling on Dirac-3 as larger problems become feasible.

## Conclusions and Outlooks

This work shows that entropy-based optimization on Dirac-3 provides a practical and effective approach for fixed-backbone computational protein design. On benchmark problems with known optima, Dirac-3 consistently achieves high-quality solutions, typically within one to two percent of CFN-derived GMEC energies. These results validate that the Hamiltonian formulation and continuous-variable optimization paradigm are well suited to realistic CPD energy landscapes.

A central finding is the favorable runtime behavior of Dirac-3. For problems within current device capacity, runtimes grow only mildly with problem size, in contrast to the rapidly escalating cost of exact classical optimization. This indicates that entropy computing may offer a scalable alternative for CPD instances where classical solvers become time-prohibitive.

For the largest proteins studied, a graph-partitioning workflow was used as a pragmatic means to fit problems within present hardware limits. This strategy enables solution of multi-thousand-variable instances but should be viewed as an interim technique rather than a core requirement. Overall, these findings establish entropy quantum computing as a promising and practical tool for large-scale computational protein design.

## Acknowledgements

The authors thank Thomas Schiex and David Simoncini for helpful discussions. Additionally, the authors gratefully acknowledge the generous support provided to this project by the Sanders Tri-Institutional Therapeutics Discovery Institute (TDI), a 509(a)(3) organization. TDI receives financial support from Takeda Pharmaceutical Company, TDI’s parent institutes (Memorial Sloan Kettering Cancer Center, The Rockefeller University, and Weill Cornell Medicine), and from a generous contribution from Mr. Lewis Sanders and other philanthropic sources.

## Methods

The decomposition approach for solving large-size protein problems is based on a graph partitioning. For each pair of positions *i* and *j* on the protein backbone, we define

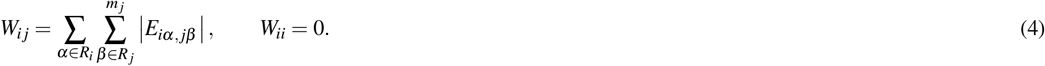

where, *E*_*iα, jβ*_ is the interaction energy between rotamer *α* at position *i*, and rotamer *β* at position *j*. The undirected, weighted graph is then *G* = (*V, E, w*) with *V* = {1, 2, …, *N*}, *E* = {{*i, j*} |*W*_*ij*_ *>* 0} and *w*(*i, j*) = *W*_*ij*_; its adjacency is *A* = [*W*_*ij*_]. Intuitively, *W*_*ij*_ summarizes the total “strength” of interactions between backbone positions *i* and *j*. Positions with large *W*_*ij*_ should preferentially remain in the same block during partitioning.

Given *G*, we seek a *k*-way partition {*S*_1_, …, *S*_*k*_} minimizing weighted cut while maintaining near-equal block sizes. We have,

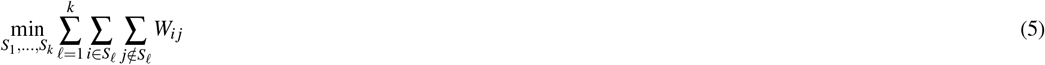

subject to,

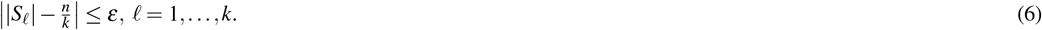

We use *k* to target hardware-fitting sub-problems (variable count of less than 953), and *ε* to control allowable load imbalance. We employ METIS, a fast, high-quality multilevel graph partitioner that scales roughly with the number of edges^57, 58^.

METIS proceeds in three stages: (1) coarsening, (2) initial *k*-way partition on a small graph, and (3) refinement. During coarsening, nodes are matched via heavy-edge matching and collapsed to produce a hierarchy *G*_0_ →*G*_1_ → · · · → G_L_ of progressively smaller graphs while approximately preserving cut structure. An initial *k*-way partition is computed on the smallest graph *G*_*L*_ (via greedy and/or KL/FM-style heuristics). The solution is then projected back up the hierarchy; at each uncoarsening level, local refinement reduces edge cut subject to balance constraints. METIS supports weighted graphs and user-specified imbalance tolerance, which is used to respect variability in *m*_*i*_ and keep sub-problems within device limits^58, 59^.

The construction of the original problem’s solution based on the subproblems is as follows. Let **x** ∈ ℝ^*n*^ denote the vectorization of all rotamer variables {*x*_*i,r*_} (flattened over positions and rotamers), and let the Dirac-3 objective function be **x**^*T*^*J***x** + *C*^*T*^**x** as was constructed above. For large proteins we partition the position graph *G* = (*V, E, w*) into *k* balanced blocks 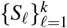. Each block *S*_*ℓ*_ induces an index set *P* ⊂ {1, …, *n*} where *n* is the total number of variables; its complement is *Q* = {1, …, *n*}*\ P*. Writing 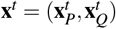 and block-partitioning *J* and *C* accordingly, the sub-problem associated with the index set *P*, and thus partition *S*_*ℓ*_, is

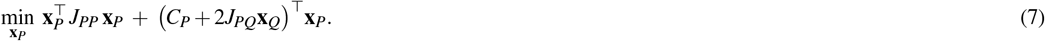

The algorithm then works as follows,

- Construct the position graph and partition it into *S*_1_, …, *S*_*k*_.
- Loop through partitions; for each partition, fix **x**_*Q*_ and solve equation 7 to get **x**_*P*_. Update **x** with values of **x**_*P*_.
- Repeat the previous step until the global solution energy change is less than a threshold.

## References

1. Carter, P. J. Introduction to current and future protein therapeutics: A protein engineering perspective. Exp. Cell Res. 317, 1261–1269, DOI: 10.1016/j.yexcr.2011.02.013 (2011).

2. Kiss, G., Celebi-Olcum, N., Moretti, R., Baker, D. & Houk, K. N. Computational Enzyme Design. Angewandte Chemie Int. Ed. 52, 5700–5725, DOI: 10.1002/anie.201204077 (2013).

3. Lutz, S. Beyond directed evolution—semi-rational protein engineering and design. Curr. Opin. Biotechnol. 21, 734–743, DOI: 10.1016/j.copbio.2010.08.011 (2010).

4. Liu, R. et al. Advances in protein engineering and its application in synthetic biology. In New Frontiers and Applications of Synthetic Biology, 147–158, DOI: 10.1016/B978-0-12-822830-4.00008-0 (Academic Press, 2022).

5. Kuhlman, B. & Bradley, P. Advances in protein structure prediction and design. Nature Reviews Molecular Cell Biology 20, 681–697, DOI: 10.1038/s41580-019-0144-x (2019).

6. Pierce, N. A. & Winfree, E. Protein design is np-hard. Protein Eng. Des. Sel. 15, 779–782 (2002).

7. Traoré, S. et al. Protein design using cost function networks. In International Conference on Principles and Practice of Constraint Programming, 787–803 (Springer, 2013).

8. Borrero, J. S., Gillen, C. & Prokopyev, O. A. A survey on quantum computing for optimization. Comput. & Oper. Res. 144, 105809 (2022).

9. Outeiral, C. et al. The prospects of quantum computing in computational molecular biology. WIREs Comput. Mol. Sci. 11, e1481 (2021).

10. Glover, Fred and Kochenberger, Gary and Du, Ying. A tutorial on formulating and using QUBO models.

11. Brubaker, J., Booth, K., Arakawa, A. et al. Quadratic unconstrained binary optimization and constraint programming approaches for lattice-based cyclic peptide docking. Sci. Reports 15, 20395 (2025).

12. Robert, A., Barkoutsos, P., Woerner, S. & Tavernelli, I. Resource-efficient quantum algorithm for protein folding. npj Quantum Inf. 7, 38 (2021).

13. Li, W. et al. A hybrid quantum computing pipeline for real-world drug discovery. Sci. Reports 14, 16942 (2024).

14. Bowie, James U and Lüthy, Roland and Eisenberg, David. A method to identify protein sequences that fold into a known three-dimensional structure. Science 253, 164–170, DOI: 10.1126/science.1853201 (1991).

15. Lippow, Shaun M and Tidor, Bruce. Progress in computational protein design. Current Opinion in Biotechnology 18, 305–311, DOI: 10.1016/j.copbio.2007.04.009 (2007).

16. Bouchiba, Younes and Cortés, Juan and Schiex, Thomas and Barbe, Sophie. Molecular flexibility in computational protein design: an algorithmic perspective. Protein Engineering, Design and Selection 34, gzab011, DOI: 10.1093/protein/gzab011 (2021).

17. Kortemme, T., Joachimiak, A., Bullock, A. N. & Baker, D. An efficient protocol for computationally designing proteins with multiple functional sites. Nat. Struct. Biol. 10, 45–52 (2003).

18. Huang, P.-S., Boyken, S. E. & Baker, D. The coming of age of de novo protein design. Nature 537, 320–327 (2016).

19. Dill, Ken A and Ozkan, S Banu and Weikl, Thomas P and Chodera, John D and Voelz, Vincent A. The protein folding problem. Annual Review of Biophysics 36, 49–69 (2007).

20. Bryngelson, Joseph D and Onuchic, José N and Socci, Nicholas D and Wolynes, Peter G. Funnels, pathways, and the energy landscape of protein folding: A synthesis. Proteins: Structure, Function, and Bioinformatics 21, 167–195 (1995).

21. Lindorff-Larsen, Kresten and Piana, Stefano and Dror, Ron O and Shaw, David E. How fast-folding proteins fold. Science 334, 517–520 (2011).

22. Jumper, John and Evans, Richard and Pritzel, Alexander and Green, Tim and Figurnov, Michael and Ronneberger, Olaf and Tunyasuvunakool, Kathryn and Bates, Russ and Žídek, Augustin and Potapenko, Anna and Ingraham, Alex and Agiomyrgiannakis, Gabriel M and Barth, Olga and Petersen, Andrew and Reiman, Stanislaw and Wifling, Erich and Jean-Baptiste, Matthew and Audrey, Helen and Chung, Lee and Green, Charlie and Homan, Thomas and Petrides, Stig and Senior, Andrew W and Kavukcuoglu, Koray and Birney, Ewan and Hassabis, Demis and Jumper, John and Hassabis, Demis. Highly accurate protein structure prediction with AlphaFold. Nature 596, 583–589 (2021).

23. Wohlwend, Jeremy and Corso, Gabriele and Passaro, Saro and Reveiz, Mateo and Leidal, Ken and Swiderski, Wojtek and Portnoi, Tally and Chinn, Itamar and Silterra, Jacob and Jaakkola, Tommi and Barzilay, Regina. Boltz-1: Democratizing Biomolecular Interaction Modeling. bioRxiv DOI: 10.1101/2024.11.19.624167 (2024).

24. Chai Discovery team and Boitreaud, Jacques and Dent, Jack and McPartlon, Matthew and Meier, Joshua and Reis, Vinicius and Rogozhonikov, Alex and Wu, Kevin. Chai-1: Decoding the molecular interactions of life. bioRxiv DOI: 10.1101/2024.10.10.615955 (2024).

25. Ponder, Jay W and Case, David A. Force fields for protein simulations. Advances in Protein Chemistry 66, 27–85 (2003).

26. Dunbrack, Roland L Jr and Karplus, Martin. Backbone-dependent rotamer library for proteins. Application to side-chain prediction. Journal of Molecular Biology 230, 543–574, DOI: 10.1006/jmbi.1993.1170 (1993).

27. Sinha, Prabhakant and Zoltners, Andris A. The Multiple-Choice Knapsack Problem. Operations Research 27, 503–515, DOI: 10.1287/opre.27.3.503 (1979).

28. Pisinger, David. A minimal algorithm for the multiple-choice knapsack problem. European Journal of Operational Research 83, 394–410 (1995).

29. Kellerer, Hans and Pferschy, Ulrich and Pisinger, David. The Multiple-Choice Knapsack Problem. In Knapsack Problems, 301–328, DOI: 10.1007/978-3-540-24777-7_11 (Springer, Berlin, Heidelberg, 2004).

30. Kuhlman, Brian and Bradley, Phil and Baker, David. Molecular recognition, design and assembly of protein structures. Current Opinion in Structural Biology 13, 435–440 (2003).

31. Kuhlman, Brian and Dantas, Gautam and Iyer, G V and Wedemeyer, William J and Smith, Leonard and Kim, Richard and Baker, David. Design of a novel globular protein fold with atomic-level accuracy. Science 302, 1388–1391 (2003).

32. Liu, Yihong and Kuhlman, Brian. Sequence design using a continuum solvent model and a two-stage hierarchical search method. Journal of Molecular Biology 358, 287–297 (2006).

33. Dahiyat, Bassil I and Mayo, Stephen L. De novo protein design: fully automated sequence selection. Science 278, 82–87, DOI: 10.1126/science.278.5335.82 (1997).

34. Leach, A. R. & Lemon, A. P. Exploring the conformational space of protein side chains using dead-end elimination and the a* algorithm. Proteins: Struct. Funct. Bioinforma. 33, 227–239 (1998).

35. Looger, Loren L and Hellinga, Homme W. Generalized dead-end elimination algorithms make large-scale protein side-chain structure prediction tractable: implications for protein design and structural genomics. Journal of Molecular Biology 307, 429–445, DOI: 10.1006/jmbi.2000.4491 (2001).

36. Lippow, Shaun M and Wittrup, K Dane and Tidor, Bruce. Computational design of antibody-affinity improvement beyond in vivo maturation. Nature Biotechnology 25, 1171–1176, DOI: 10.1038/nbt1332 (2007).

37. Hallen, M. A., Keedy, D. A. & Donald, B. R. Dead-end elimination with perturbations (deeper): a provably accurate algorithm for protein design. J. Comput. Biol. 20, 740–761 (2013).

38. Desmet, J., De Maeyer, M., Hazes, B. & Lasters, I. The dead-end elimination theorem and its use in protein side-chain positioning. Nature 356, 539–542 (1992).

39. Loos, B., Fackeldey, K. & Schomburg, D. Dead-end elimination with backbone flexibility. J. Mol. Biol. 326, 1–9 (2003).

40. Leaver-Fay, A. et al. Rosetta3: an object-oriented software suite for the simulation and design of macromolecules. Methods Enzymol. 487, 545–574 (2011).

41. Simoncini, David and Allouche, David and de Givry, Simon and Katsirelos, George and Schiex, Thomas and Barbe, Sophie. Guaranteed Discrete Energy Optimization on Large Protein Design Problems. Journal of Chemical Theory and Computation 11, 5980–5989, DOI: 10.1021/acs.jctc.5b00594 (2015).

42. Allouche, David and de Givry, Simon and Katsirelos, George and Schiex, Thomas and Traoré, Seydou and Barbe, Sophie. Computational protein design as an optimization problem. Artificial Intelligence 212, 100–115, DOI: 10.1016/j.artint.2014.03.003 (2014).

43. Kingsford, C. L., Chazelle, B. & Singh, M. Solving and analyzing side-chain positioning problems using linear and integer programming. Bioinformatics 21, 1028–1036 (2005).

44. Alford, R. F. et al. The rosetta all-atom energy function for macromolecular modeling and design. J. Chem. Theory Comput. 13, 3031–3048 (2017).

45. Gainza, P., Roberts, K. E. & Donald, B. R. Osprey: protein design with ensembles, flexibility, and provable algorithms. Methods Enzymol. 523, 87–107 (2013).

46. Allouche, D., de Givry, S., Katsirelos, G., Schiex, T. & Zytnicki, M. Anytime hybrid best-first search with tree decomposition for weighted csp. In Proceedings of the 16th International Conference on Principles and Practice of Constraint Programming, 12–27 (Springer, 2010).

47. Xu, J. & Berger, B. Protein design using fragment assembly and branch-and-bound search. Proc. Natl. Acad. Sci. 102, 9753–9758 (2005).

48. Perdomo-Ortiz, A., Dickson, N., Drew-Brook, M., Rose, G. & Aspuru-Guzik, A. Finding low-energy conformations of lattice protein models by quantum annealing. Sci. Reports 2, 571 (2012).

49. Babbush, R., Perdomo-Ortiz, A., O’Gorman, B., Macready, W. G. & Aspuru-Guzik, A. Adiabatic quantum simulation of protein folding. arXiv preprint arXiv:1211.3422 (2014).

50. Nguyen, L. et al. Entropy computing, a paradigm for optimization in open photonic systems. Commun. Phys. 8, 411 (2025).

51. Chukwu, U., Miri, M.-A. & Chancellor, N. Explicitly quantum-parallel computation by displacements (2025). 2510.19730.

52. Emami, B. et al. Financial fraud detection with entropy computing. arXiv preprint arXiv:2503.11273 (2025).

53. Rohl, C. A., Strauss, C. E., Misura, K. M. & Baker, D. Protein structure prediction using rosetta. Methods Enzymol. 383, 66–93 (2004).

54. Rubenstein, A. B., Blacklock, K., Nguyen, H., Case, D. A. & Khare, S. D. Systematic comparison of amber and rosetta energy functions for protein structure evaluation. J. Chem. Theory Comput. 14, 6015–6025 (2018).

55. Allouche, D. et al. Computational protein design as an optimization problem. Artif. Intell. 212, 59–79 (2014).

56. de Givry, S., Schiex, T. & the toulbar2 team. toulbar2: An open-source exact solver for cost function networks. https://toulbar2.github.io/toulbar2/ (2023). Accessed 2025-09-22.

57. Karypis, G. & Kumar, V. A fast and high quality multilevel scheme for partitioning irregular graphs. SIAM J. on Sci. Comput. (1998). Also available as a widely cited technical report; see METIS materials.

58. Karypis, G. METIS 5.1.x Manual. Karypis Lab (2013). User’s guide and API for METIS 5.x.

59. Karypis, G. & Kumar, V. Multilevel k-way partitioning scheme for irregular graphs. J. Parallel Distributed Comput. 48, 96–129 (1998).

