## Supplementary Information for "Entropy Quantum Computing for Fixed-Backbone Protein Design"

### The fixed-backbone approach using a rotamer library

Table S1 details the 20 natural amino acids and their corresponding rotamer counts derived from the Dunbrack-Karplus library used in our protein design formulation[1]. These values define the state space for each position in the protein backbone, directly influencing the dimensionality and complexity of the resulting quadratic Hamiltonian. The significant variance in rotamer counts, ranging from a single state for Glycine to 243 for Arginine, highlights the non-uniform combinatorial challenges. Table S2 reports problem sizes and coefficient dynamic ranges for all of the proteins studies in this paper.

**Table S1.** The names, three-letter codes, and one-letter codes of the 20 natural amino acids along with the number of rotamers in the Dunbrack-Karplus rotamer library

| Amino Acid Name | Three-Letter Code | One-Letter Code | Number of Rotamers In Dunbrack Karplus Library |
|---|---|---|---|
| Alanine | ALA | A | 1 |
| Arginine | ARG | R | 243 |
| Asparagine | ASN | N | 9 |
| Aspartic Acid | ASP | D | 9 |
| Cysteine | CYS | C | 3 |
| Glutamic Acid | GLU | E | 27 |
| Glutamine | GLN | Q | 27 |
| Glycine | GLY | G | 1 |
| Histidine | HIS | H | 9 |
| Isoleucine | ILE | I | 3 |
| Leucine | LEU | L | 9 |
| Lysine | LYS | K | 81 |
| Methionine | MET | M | 27 |
| Phenylalanine | PHE | F | 9 |
| Proline | PRO | P | 3 |
| Serine | SER | S | 3 |
| Threonine | THR | T | 3 |
| Tryptophan | TRP | W | 9 |
| Tyrosine | TYR | Y | 9 |
| Valine | VAL | V | 3 |

| Protein | Size | Dynamic Range (dB) |
|---|---|---|
| 1MJC | 493 | 49.35 |
| 1CSK | 616 | 49.36 |
| 1SHF | 638 | 49.39 |
| 1SHG | 737 | 49.39 |
| 1NXB | 800 | 49.48 |
| 1TEN | 808 | 49.45 |
| 1POH | 943 | 49.54 |
| 1RIS | 3276 | 49.63 |
| 1GVP | 3826 | 49.66 |

**Table S2.** Mid-sized and large-sized CPD benchmarks: variable counts and coefficient dynamic ranges of the Hamiltonians.

### The Dirac-3 Entropy Quantum Computing Device

Quantum Computing Inc.'s Dirac-3 is a third-generation entropy-based quantum-inspired optimization device designed for solving large-scale, optimization problems in science and industry. Unlike gate-model quantum computers or quantum annealers, which rely respectively on unitary circuit operations or simulated Boltzmann sampling, Dirac-3 implements an Entropy Quantum Computing (EQC) paradigm in which optimization is performed by minimizing Hamiltonians within an open quantum system[2]. The device uses photonic and optical components to encode problem Hamiltonians directly and then applies entropy gradients to drive the system toward low-energy configurations.

A key feature of Dirac-3 is its ability to handle Hamiltonians with very wide dynamic ranges (in practice up to 40–70 dB), meaning that both weak and strong interactions can be represented simultaneously without rescaling. This is particularly important in protein design, where interaction energies may differ by orders of magnitude (for example, between steric clashes, hydrogen bonding, and electrostatic interactions). By maintaining high fidelity across these ranges, Dirac-3 avoids loss of resolution that would otherwise compromise solution quality.

Another distinctive advantage is the device's global connectivity. Many quantum annealers require embedding logical variables into sparse hardware graphs, which introduces overhead and noise. In contrast, Dirac-3 provides all-to-all connectivity, enabling dense interaction graphs such as those arising in CPD to be encoded natively. This connectivity is central for modeling pairwise rotamer interactions without approximation.

From a practical standpoint, Dirac-3 is engineered for low-power operation (approximately 80–100 W) and supports user access through QCi's software interface, which allows Hamiltonians to be specified in terms of linear biases and quadratic (or higher-order) couplings. The software stack also enables hyper-parameter control (e.g., relaxation schedules, sample counts, and dynamic range parameters) that tailor the device's performance to the problem instance.

Beyond protein design, Dirac-3 has already been demonstrated in several domains, including financial portfolio optimization, supply chain logistics, and anomaly detection. For example, recent work in fraud detection benchmarked Dirac-3 against classical machine learning models and showed competitive performance while maintaining favorable scaling as the dimensionality of the Hamiltonian increased[3]. These demonstrations highlight Dirac-3 as a flexible, domain-agnostic optimization platform, motivating its exploration for computational protein design in the present study.

### Device parameters and hyper-parameters

The following tables summarize results on the impact of device parameters and hyper-parameters. Table S3 shows the best energy found for the protein 1MJC and the average device runtime when considering different mean photon numbers. Table S4 shows the best energy and run-time for the protein 1MJC over different relaxation schedules of Dirac-3. In Table S5, max-thresholds in the 5–10 range yield better solution energies by altering the Hamiltonian just enough to keep the dynamic range manageable; pushing to 100 degrades solution energy due to larger dynamic range values. In Table S6, min-thresholds around $10^{-4}$ to $10^{-3}$ are conservative and stable; while $10^{-1}$ happens to give a slightly lower energy here, that behavior is instance-specific and risks over-pruning. Tables S7 and S8 show the impact of the penalty multipliers $\alpha$ and $\beta$.

| Mean Photon Number | Best Dirac-3 Energy | Average Sample Runtime (s) |
|---|---|---|
| 0.000066 | -70.03 | 40.35 |
| 0.0001 | -70.17 | 27.14 |
| 0.001 | -70.45 | 3.29 |
| 0.002 | -70.45 | 1.96 |
| 0.003 | -70.45 | 2.10 |
| 0.004 | -70.20 | 1.43 |
| 0.005 | -70.18 | 1.28 |
| 0.0066 | -69.48 | 1.14 |

**Table S3.** Solution energy and device runtime versus mean photon number for 1MJC protein; relaxation schedule: 1; num samples = 100; max energy threshold = 10; min energy threshold = 0.001; $\alpha = 20$; $\beta = 0$.

| Relaxation Schedule | Best Dirac-3 Energy | Average Sample Runtime (s) |
|---|---|---|
| 1 | -70.28 | 2.12 |
| 2 | -70.45 | 4.23 |
| 3 | -70.35 | 6.22 |
| 4 | -70.35 | 12.78 |

**Table S4.** Solution energy and device runtime versus relaxation schedule for 1MJC protein; mean photon number: 0.003; num samples = 100; max energy threshold = 10; min energy threshold = 0.001; $\alpha = 20$; $\beta = 0$.

| Max Energy Threshold | Dynamic Range | Best Dirac-3 Energy |
|---|---|---|
| 1 | 49.35 | -70.54 |
| 5 | 49.35 | -70.45 |
| 10 | 49.35 | -70.40 |
| 50 | 49.35 | -69.57 |
| 100 | 50.55 | -66.01 |

**Table S5.** Solution energy and dynamic range versus max energy threshold for 1MJC protein; relaxation schedule = 2; mean photon number: 0.003; num samples = 100; min energy threshold = 0.001; $\alpha = 20$; $\beta = 0$.

| Min Energy Threshold | Dynamic Range | Best Dirac-3 Energy |
|---|---|---|
| $1.0 \times 10^{-6}$ | 79.35 | -70.45 |
| $1.0 \times 10^{-5}$ | 79.35 | -70.33 |
| $1.0 \times 10^{-4}$ | 59.35 | -70.45 |
| $1.0 \times 10^{-3}$ | 49.35 | -70.40 |
| $1.0 \times 10^{-2}$ | 39.35 | -70.40 |
| $1.0 \times 10^{-1}$ | 29.35 | -70.35 |

**Table S6.** Solution energy and dynamic range versus min energy threshold for 1MJC protein; relaxation schedule = 2; mean photon number: 0.003; num samples = 100; max energy threshold = 10; $\alpha = 20$; $\beta = 0$.

| $\alpha$ | Dynamic Range | Best Dirac-3 Energy |
|---|---|---|
| 5.0 | 44.16 | -70.45 |
| 10.0 | 46.63 | -70.45 |
| 20.0 | 49.35 | -70.40 |
| 50.0 | 53.15 | -70.47 |
| 100.0 | 56.09 | -69.88 |

**Table S7.** Solution energy and dynamic range versus $\alpha$ for 1MJC protein; relaxation schedule = 2; mean photon number: 0.003; num samples = 100; max energy threshold = 10; min energy threshold = 0.001; $\beta = 0$.

| $\beta$ | Dynamic Range | Dirac-3 Energy |
|---|---|---|
| 0.0 | 49.35 | -70.40 |
| 0.1 | 49.14 | -70.45 |
| 0.5 | 48.20 | -70.42 |

**Table S8.** Solution energy and dynamic range versus $\alpha$ for 1MJC protein; relaxation schedule = 2; mean photon number: 0.003; num samples = 100; max energy threshold = 10; min energy threshold = 0.001; $\alpha$ = 20.

### References


1. Dunbrack, Roland L Jr and Karplus, Martin. Backbone-dependent rotamer library for proteins. Application to side-chain prediction. *Journal of Molecular Biology* **230**, 543–574, DOI: 10.1006/jmbi.1993.1170 (1993).
2. Nguyen, L. *et al.* Entropy computing, a paradigm for optimization in open photonic systems. *Commun. Phys.* **8**, 411 (2025).
3. Emami, B. *et al.* Financial fraud detection with entropy computing. *arXiv preprint arXiv:2503.11273* (2025).
